# DoubletFinder: Doublet detection in single-cell RNA sequencing data using artificial nearest neighbors

**DOI:** 10.1101/352484

**Authors:** Christopher S. McGinnis, Lyndsay M. Murrow, Zev J. Gartner

## Abstract

Single-cell RNA sequencing (scRNA-seq) using droplet microfluidics occasionally produces transcriptome data representing more than one cell. These technical artifacts are caused by cell doublets formed during cell capture and occur at a frequency proportional to the total number of sequenced cells. The presence of doublets can lead to spurious biological conclusions, which justifies the practice of sequencing fewer cells to limit doublet formation rates. Here, we present a computational doublet detection tool – DoubletFinder – that identifies doublets based solely on gene expression features. DoubletFinder infers the putative gene expression profile of real doublets by generating artificial doublets from existing scRNA-seq data. Neighborhood detection in gene expression space then identifies sequenced cells with increased probability of being doublets based on their proximity to artificial doublets. DoubletFinder robustly identifies doublets across scRNA-seq datasets with variable numbers of cells and sequencing depth, and predicts false-negative and false-positive doublets defined using conventional barcoding approaches. We anticipate that DoubletFinder will aid in scRNA-seq data analysis and will increase the throughput and accuracy of scRNA-seq experiments.

## INTRODUCTION

Since its introduction nearly a decade ago, scRNA-seq has been used to elucidate previously unknown cell types and reconstruct developmental dynamics among heterogeneous cell populations (Human Cell Atlas Consortium, 2017). At first, scRNA-seq workflows were limited to tens to hundreds of cells which hindered data interpretation due to batch effects and low statistical power (Stegle et al., 2016). Today, sequencing thousands to hundreds of thousands of cells is routine due to the advent of droplet microfluidics and nanowell-based sequencing strategies (Macosko et al., 2015; Klein et al., 2015; Zheng et al., 2017; Gierahn et al., 2017; Takara Bio USA, 2018). These techniques rely on a Poisson loading strategy to compartmentalize individual cells and mRNA capture beads before cell lysis, mRNA capture, and transcript barcoding via reverse transcription. Since cells are captured randomly, the proportion of droplets containing >1 cell – known as doublets – scales linearly across an experimentally-relevant range of input cell concentrations (10X Genomics, 2017), justifying the practice of limiting the number of sequenced cells to minimize doublet formation rates.

The confounding effects of doublets in scRNA-seq data are well-appreciated (Ilicic et al., 2016). However, genomic and cellular barcoding techniques for identifying doublets have only recently been developed (Stoeckius et al., 2017; Kang et al., 2018; Gehring et al., 2018; Guo et al., 2018; Rosenberg et al., 2018). In one such strategy, distinct samples receive unique oligonucleotide barcodes delivered by conjugation to antibodies targeting broadly expressed cell-surface antigens. When the barcoded pools are combined and sequenced, doublets can be identified according to the co-occurrence of orthogonal cell ‘hashtags’ (Stoeckius et al., 2017). In a second strategy, doublets in a pooled population of cells from different individuals are identified by a computational pipeline, Demuxlet, which facilitates doublet inference based on the co-occurrence of mutually-exclusive SNP profiles (Kang et al., 2018).

By detecting doublets, both Demuxlet and Cell Hashing minimize technical artifacts while enabling users to “superload” droplet microfluidics devices for increased scRNA-seq throughput. However, both methods have limitations. First, neither method can identify doublets formed from identically-barcoded cells. Second, neither method is universally applicable across experimental systems, since Demuxlet requires genetically distinct samples and Cell Hashing requires unique antibody-oligonucleotide conjugate panels for the cell types and species of interest. Third, neither method can be used to analyze existing scRNA-seq datasets. For these reasons, computational methods for defining doublets based on gene expression patterns alone are highly desirable.

Here, we present DoubletFinder, a computational doublet detection tool that relies solely on gene expression data. Beginning with the observation that doublets cluster separately from singlets in high-dimensional gene expression space (Stoeckius et al., 2017; Kang et al., 2018), we reasoned that real doublets would cluster together with synthetic doublets formed by averaging the expression data of two real cells. By merging artificial doublets with existing scRNA-seq data, we can distinguish doublets from singlets according to the proportion of artificial nearest neighbors (pANN) for each real cell in gene expression space. Thresholding the resulting pANN distribution to match the expected number of doublets provides an accurate metric for doublet prediction that can be applied to any scRNA-seq dataset.

## RESULTS

### DoubletFinder predicts doublets more accurately than nUMIs

Existing strategies for identifying doublets using gene expression features primarily rely on two sources of information. First, since the total number of captured mRNA molecules is expected to be greater for doublets than singlets, doublets are commonly excluded by thresholding cells with high numbers of unique molecular identifiers (nUMIs; Islam et al., 2014; Ziegenhain et al., 2017). While intuitively appealing, technical variability in mRNA capture efficiency and biological variability in mRNA content limits the utility of nUMI-based doublet predictions (Stoeckius et al., 2017). In a second strategy, doublets are removed from scRNA-seq data by identifying groups of cells exhibiting the co-expression of genes with non-overlapping expression patterns *in vivo* (Rosenberg et al., 2018). This strategy cannot be applied to biological systems where such marker genes are unknown, undetected, or unavailable. Moreover, such a strategy could theoretically lead to the erroneous removal of new cell types or developmental states with intermediate expression profiles (Fig. 1A). Given these shortcomings, new methods for predicting doublets using gene expression features alone would greatly benefit the single-cell genomics field.

**Figure 1:**
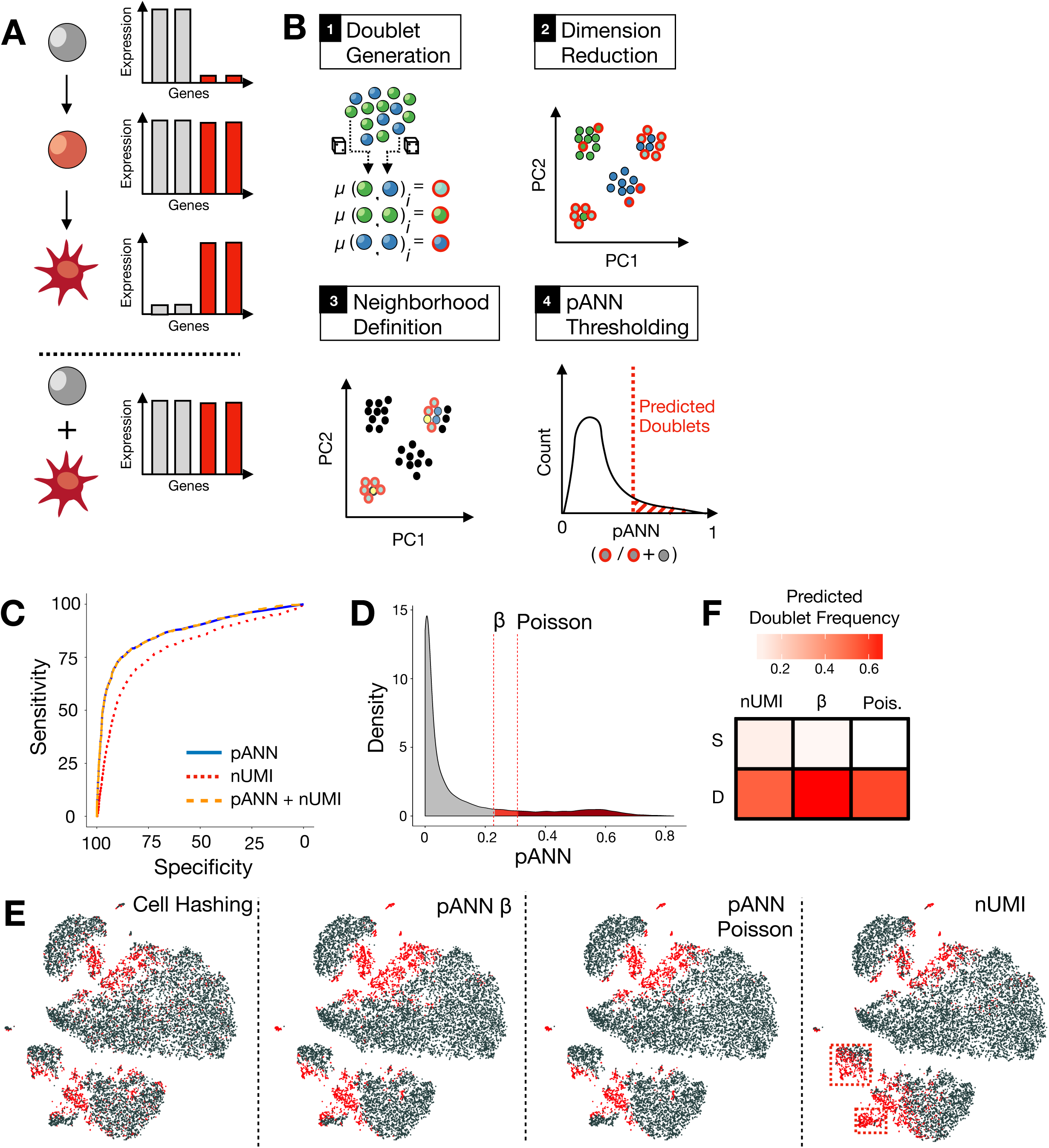
DoubletFinder robustly predicts Cell Hashing doublets and outperforms nUMI thresholding. (A) Schematic describing importance of doublet detection. Developmental intermediates (light red) can express genes associated with both progenitor (grey) and differentiated (dark red) cell types. Doublets formed from progenitor and mature cells may mimic the expression profile of intermediate cell states, and thereby confound analysis. (B) Schematic of DoubletFinder workflow. After artificial doublet (red outline) generation, the proportion of artificial nearest neighbors is defined for every real cell (examples highlighted yellow). These results are thresholded to define doublet predictions. (C) Density plot of pANN values with red dotted lines denoting expected doublet thresholds. Histogram is colored according to whether the cells were called as singlets (grey) or doublets by one (light red) or both thresholding strategies (dark red). (D) t-SNE visualization of real and predicted Cell Hashing doublets (red) and singlets (grey), DoubletFinder predictions, and nUMI thresholding predictions. nUMI predictions deviate significantly from Cell Hashing results (red dashed boxes). (E) Heat map showing the proportion of DoubletFinder and nUMI-predicted doublets present in Cell Hashing singlet (S) and doublet (D) groups. (F) ROC analysis of logistic regression models trained using nUMIs alone (dotted red), pANN alone (solid blue) and both nUMIs and pANN (dashed orange) as features.

DoubletFinder predicts doublets in a fashion agnostic to nUMIs, marker gene expression, genetic background or exogenous barcodes and can be split into four distinct steps: (1) Generate artificial doublets, (2) Merge real and artificial data and reduce dimensionality with principal component analysis (PCA), (3) Define the nearest neighbors for every real cell in PC space, and (4) Compute and threshold the proportion of artificial nearest neighbors (pANN; Fig. 1B). We tested the efficacy of DoubletFinder against scRNA-seq datasets where doublets are empirically-defined: The publically-available Cell Hashing and Demuxlet datasets comprised of 15,178 and 35,524 peripheral blood mononuclear cells (PBMCs), respectively. Demuxlet PBMCs were derived from 8 genetically-distinct human sources while Cell Hashing PBMCs were barcoded with 8 distinct antibody-oligonucleotide conjugate panels. Using optimized input parameters (Supplementary Materials, Fig. S1), we tested whether pANN outperforms nUMIs as a doublet-prediction feature by using receiver operating curve (ROC) analysis to compare logistic regression models trained on the Cell Hashing data using pANN alone, nUMIs alone, or both (Fig. 1C). ROC analysis demonstrates that pANN predicts doublets more accurately than nUMIs. Moreover, the model trained with both features performed nearly indistinguishably to the pANN-alone model, suggesting that DoubletFinder captures all of the doublet-specific information inherent to nUMIs.

### DoubletFinder predicts Cell Hashing doublets

To make specific doublet predictions for each cell, DoubletFinder rank-orders cells by their pANN values and thresholds this list according to the number of expected doublets. To test the robustness of pANN thresholding, the number of expected doublets was determined using two different strategies (Fig. 1D). First, since 8 samples were multiplexed in the Cell Hashing and Demuxlet studies, we reasoned that 1/8 of the true doublets were undetected because they were formed from genetically-identical or identically-barcoded cells. Thus, we thresholded pANN according to the number of detected doublets with an assumed 12.5% false negative rate (b). Second, since the doublet formation rate can be accurately estimated by applying Poisson statistics to the number of cells loaded into the droplet microfluidics device (10X Genomics, 2017), we thresholded pANN according to this rate. For standard scRNA-seq experiments where doublets are not empirically-defined, pANN can only be thresholded using the Poisson strategy.

Depending on the threshold used, DoubletFinder predicted 2680 or 2155 doublets when applied to the full Cell Hashing dataset. Single-cell gene expression data was visualized using t-stochastic neighborhood embedding (t-SNE; van der Maaten and Hinton, 2008) and cells were colored according to their real and predicted doublet status (Fig. 1E). Visual comparison of doublets in t-SNE space illustrates that DoubletFinder predictions closely track Cell Hashing results. This result is further supported by the observation that the frequency of DoubletFinder predictions is highly enriched in Cell Hashing-defined doublet groups relative to singlet groups (Fig. 1F), regardless of the thresholding strategy used.

### DoubletFinder predicts Cell Hashing false-negatives

DoubletFinder predictions exhibit a less ‘speckled’ appearance in t-SNE space relative to the Cell Hashing results (Fig. 2A, insets). Considering that doublets often cluster separately from singlets in gene expression space (Stoeckius et al., 2017; Kang et al., 2018), we reasoned that cells called as singlets via Cell Hashing, that nonetheless co-cluster with high-confidence doublets, are actually false-negatives derived from identically-barcoded cells. Two main predictions follow from this line of reasoning. First, if the putative false negatives are truly doublets, then they should exhibit gene expression patterns associated with distinct cell types. In line with this prediction, Cell Hashing-defined doublets and singlets in the highlighted region express marker genes for both B cells and NK cells (Fig. 2B) – hematopoietic cell types that do not share a common progenitor in peripheral blood. Second, since false negative cells would be associated with the combined barcodes of two cells, the nUMI counts for the most abundant barcode should be significantly higher in false negatives than high-confidence doublets. Moreover, since false negatives would not be associated with high levels of multiple barcodes, the second most abundant barcode should be similar to high-confidence singlets and significantly lower relative to high-confidence doublets. Statistical analysis supports these predictions (Wilcoxon rank sum test, p < 10^-13^; Fig. 2C). Collectively, these results demonstrate that DoubletFinder robustly recapitulates doublet assignments and accurately predicts Cell Hashing false-negatives.

**Figure 2:**
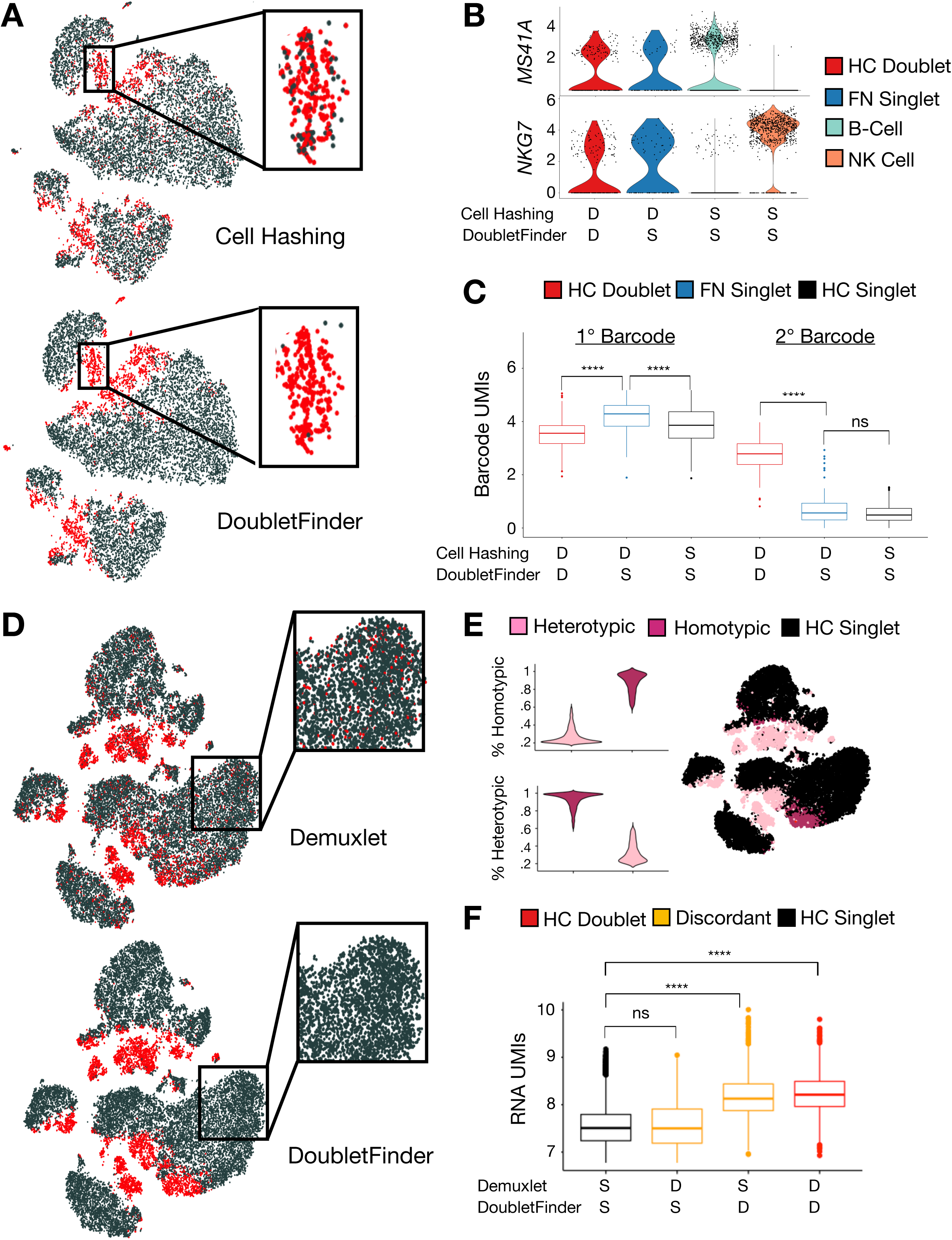
DoubletFinder detects false-negative and false-positive doublet predictions in Cell Hashing and Demuxlet datasets. (A) t-SNE visualization of real and predicted Cell Hashing doublets (red) and singlets (grey) highlighting a doublet-enriched, ‘speckled’ region. (B) Violin plots describing the distribution of marker gene expression in high-confidence doublets (red), putative false-negatives (blue), and singlet B-cells (teal) and NK cells (orange). (C) Barcode UMI box plots for the 1^st^ and 2^nd^ most abundant barcodes in high-confidence singlets (black) and doublets (red), as well as false-negative singlets (blue). (D) t-SNE visualization of real and predicted Demuxlet doublets (red) and singlets (grey) highlighting putative false-positives. (E) Violin plots describing the contributions of homotypic and heterotypic doublets to each DoubletFinder-defined doublet’s nearest neighborhood. t-SNE visualization of putative homotypic and heterotypic doublet clusters in gene expression space. (F) RNA UMI box plots for high-confidence singlets (black) and doublets (red) as well as discordant doublet predictions between Demuxlet and DoubletFinder (orange).

### DoubletFinder predicts Demuxlet doublets and identifies putative false-positives

To test whether DoubletFinder performance is sensitive to changes in the number of sequenced cells and sequencing depth, we applied DoubletFinder to the Demuxlet dataset. In addition to having more cells than the Cell Hashing data, the average number of UMIs (2408 vs 676) and genes (837 vs 376) per cell is also greater in the Demuxlet data. In line with our previous results, visual comparison of real and predicted doublets using t-SNE illustrates that DoubletFinder successfully identifies all doublet-enriched regions in gene expression space (Fig. 2D).

As with our Cell Hashing comparison, there were a number of regions in gene expression space where DoubletFinder predictions differed from Demuxlet classifications. Specifically, there were many DoubletFinder-defined doublets called as singlets by Demuxlet that give doublet-enriched clusters the ‘speckled’ appearance discussed above. Moreover, in contrast to the Cell Hashing comparison, there was a subset of cells classified as doublets by Demuxlet and singlets using DoubletFinder (Fig. 2D, insets). Interestingly, the majority of these discordant calls are scattered amongst high-confidence singlet clusters in gene expression space. This observation can be explained by two alterative models. In one model, these discordant calls are caused by homotypic doublets – i.e., doublets formed from cells of the same type – which presumably have a similar transcriptional profile to singlets and, thus, would be more difficult for DoubletFinder to detect relative to heterotypic doublets. Alternatively, the discordant calls are due to false-positive Demuxlet classifications.

If these cells were in fact homotypic doublets left undetected by DoubletFinder, then one would expect that DoubletFinder was insensitive to homotypic doublets throughout the Demuxlet dataset. To test this possibility, we tracked the cell types comprising each artificial doublet and deconvolved pANN values into homotypic and heterotypic components. Visualization of cells with majority homotypic or heterotypic nearest neighbors highlights a region of homotypic doublets formed from CD4^+^ T-cells (Fig. 2E; Supplementary Materials, Fig. S2), which suggests that DoubletFinder has the sensitivity to detected certain classes of homotypic doublets. Moreover, the scattering of discordant doublet classifications amongst high-confidence singlet clusters in gene expression space is also evident in Demuxlet classifications of the Cell Hashing data (Supplementary Materials, Fig. S2). Cell Hashing is sensitive to homotypic doublets, which suggests that the discordant calls are not a consequence of homotypic doublets missed by DoubletFinder. Finally, if the putative false-positive calls are homotypic doublets, one would expect the number of RNA UMIs to approximate levels observed for high-confidence doublets. Interestingly, the RNA nUMI distribution for putative false-positives is nearly indistinguishable to high-confidence singlets (Wilcoxon rank sum test, p = 0.34), while high-confidence doublets and putative false-negatives are both significantly enriched for RNA nUMIs (Wilcoxon rank sum test, p < 10^-15^; Fig. 2F). While it is difficult to definitively ascertain the ground truth for these discordant calls, these results collectively demonstrate that DoubletFinder is robust across a range of cell numbers and sequencing depths and prospectively predicts Demuxlet false-positives.

## DISCUSSION

High-throughput scRNA-seq suffers from the formation of doublets due to the inherent nature of Poisson cell loading. Doublets can lead to spurious conclusions during analysis when left unidentified because the resulting artefactual expression data may be interpreted as previously-undescribed cell types, developmental intermediates, or disease states. As a result, it has become common practice to minimize the doublet formation rate by minimizing the ratio of sequenced cells to mRNA capture beads. Although recent advances in direct doublet detection methodologies have proven to be effective, they are not universally or retroactively applicable. For this reason, complementary techniques for predicting doublets based only on gene expression data have the potential to further increase scRNA-seq throughput while removing technical artifacts.

Towards this goal, DoubletFinder accurately identifies doublets in scRNA-seq data by integrating artificial doublets into real data and computing the pANN for every real cell. We have shown that DoubletFinder distinguishes real doublets from singlets better than nUMIs in the Cell Hashing dataset. Moreover, we demonstrate that DoubletFinder accurately predicts doublets for two independent PBMC scRNA-seq datasets of different sizes and sequencing depths. As these are the only publically available data with empirically-defined doublets, it is unclear whether DoubletFinder will require further optimization for scRNA-seq datasets describing different tissues or biological systems. We have also shown that DoubletFinder identifies false-negative and putative false-positive doublet classifications present in these datasets, which supports the use of DoubletFinder in concert with Cell Hashing, Demuxlet, and other barcoding approaches. Finally, we demonstrate that DoubletFinder performs robustly with pANN thresholding strategies that differed by >5000 cells (Fig. 1D). This suggests that DoubletFinder can be applied in experimental contexts where doublet formation rates differ significantly from industry estimates – e.g., clumpy single-cell suspensions or especially cohesive cell types. Collectively, DoubletFinder represents a fast, easy-to-use doublet detection strategy that will aid the single-cell genomics community in data analysis and enable high-throughput scRNA-seq technologies to be utilized to their fullest potential.

## MATERIALS & METHODS

### DoubletFinder Overview

Artificial doublets were generated from raw UMI count matrices via random sampling of cell expression profiles without replacement before pre-processing using the ‘Seurat’ R package, as described previously (Butler et al., 2018). Notably, no sources of variation were regressed out of the merged data before PCA, and the top 10 PCs – chosen via inflection point estimation on the corresponding elbow plot – were used to define the Euclidean distance matrix using the ‘dist’ R function.

### ROC Analysis

ROC analysis-based model comparisons were performed using the ‘ROCR’ (Sing et al., 2005) and ‘pROC’ (Robin et al., 2011) R packages. Briefly, logistic regression models were defined on a training set comprising half the total data using the ‘glm’ R function with the link argument set to ‘logit’. These models were then used to create a vector describing each cell’s doublet probability with the ‘predict’ R function. ROC analysis was then performed by calculating the sensitivity and specificity of doublet predictions based on the aforementioned probability vector at varying probability thresholds. The AUC was then calculated for the resulting curve, and AUC was used as a proxy for doublet detection model performance.

#### Statistical Analysis

Statistically-significant differences between UMI levels were defined using the Wilcoxon rank sum test implemented with the ‘pairwisewilcox.test’ R function. Multiple comparison correction was performed using the Benjamini-Hochberg procedure.

#### Data Availability

Cell Hashing (GEO: GSE108313) and Demuxlet (GEO: GSE96583) UMI count matrices were downloaded from the Gene Expression Omnibus. DoubletFinder is implemented as a fast, easy-to-use R package that interfaces with Seurat version 2.0 and higher. DoubletFinder can be downloaded from GitHub (github.com/chris-mcginnis-ucsf/DoubletFinder).

## ACKNOWLEDGMENTS

We thank Matt Thomson (California Institute of Technology) for helpful discussion, as well as Jimmie Ye (UCSF), Marlon Stoeckius and Shiwei Zheng (New York Genome Center) for providing data access.

## AUTHOR CONTRIBUTIONS

C.S.M., L.M.M., and Z.J.G conceptualized the method and wrote the manuscript. C.S.M. wrote the software and performed all bioinformatics analyses.

